# Random forest-based modelling to detect biomarkers for prostate cancer progression

**DOI:** 10.1101/602334

**Authors:** Reka Toth, Heiko Schiffmann, Claudia Hube-Magg, Franziska Büscheck, Doris Höflmayer, Sören Weidemann, Patrick Lebok, Christoph Fraune, Sarah Minner, Thorsten Schlomm, Guido Sauter, Christoph Plass, Yassen Assenov, Ronald Simon, Jan Meiners, Clarissa Gerhäuser

## Abstract

The clinical course of prostate cancer (PCa) is highly variable, demanding an individualized approach to therapy and robust prognostic markers for treatment decisions. We present a random forest-based classification model to predict aggressive behaviour of PCa. DNA methylation changes between PCa cases with good or poor prognosis (discovery cohort with n=70) were used as input. The model was validated with data from two large independent PCa cohorts from the “International Cancer Genome Consortium” (ICGC) and “The Cancer Genome Atlas” (TCGA). Ranking of cancer progression-related DNA methylation changes allowed selection of candidate genes for additional validation by immunohistochemistry. We identified loss of ZIC2 protein expression, mediated by alterations in DNA methylation, as a promising novel prognostic biomarker for PCa in >12,000 tissue micro-array tumors. The prognostic value of ZIC2 proved to be independent from established clinico-pathological variables including Gleason grade, tumor stage, nodal stage and PSA. In summary, we have developed a PCa classification model, which either directly or *via* expression analyses of the identified top ranked candidate genes might help in decision making related to the treatment of prostate cancer patients.

## Introduction

Prostate cancer (PCa) is the second most prevailing cancer in the male population worldwide, with an estimated 1.1 Mio. newly diagnosed cases and 307.500 cancer-related deaths in 2012 (Ferlay et al, 2015; Stewart & Wild, 2014). Although the aetiology of PCa is controversial, it is likely to result from accumulating DNA damage in stress-exposed ageing prostate epithelial cells (Gelmann, 2008). Specifically, chromosomal rearrangements and oncogene fusion genes in these cells are driven by androgens (Weischenfeldt et al, 2013). Despite a large number of studies that have suggested a multitude of candidate prognostic markers in PCa (Schlomm et al, 2007), none of these genes has proven to be superior over the established histological prognostic factors including tumour stage and Gleason grade. Localized PCa with low Gleason score usually remains indolent, requiring only active surveillance or minimal treatment. Nevertheless, many patients may be over-treated with associated side effects and substantial costs (Cooperberg et al, 2011). There is, therefore, general agreement that novel specific biomarkers for the diagnosis and prognosis of PCa are needed for an efficient clinical management of this disease (Prensner et al, 2012).

Recent high-resolution genome-wide studies have significantly improved our understanding of chromosomal and genetic alterations associated with PCa development, such as the androgen-driven formation of gene fusions between the transmembrane serine protease TMPRSS2 and a member of the oncogenic ETS transcription factor family like ERG in about 50% of all PCa cases, and frequent loss of the tumour suppressor gene PTEN (Cancer Genome Atlas Research, 2015; Fraser et al, 2017; Gerhauser et al, 2018; Taylor et al, 2010; Tomlins et al, 2005; Weischenfeldt et al, 2013). These events affect signalling pathways and lead to alterations in gene expression programs that have been used for the development of gene signatures (genomic classifiers) as biomarkers for the prediction of PCa prognosis. Although several studies have demonstrated some prognostic value of gene expression-based signatures, due to the limited stability of RNA and often low quality when extracted from formalin-fixed paraffin-embedded (FFPE) material, protein- or DNA-based methods might be superior to RNA expression profiles for biomarker development.

There is substantial evidence that genetic defects in PCa are complemented or even preceded by epigenetic aberrations such as DNA methylation (Florl et al, 2004; Nelson et al, 2009). Methylation of CpG rich sequences in gene promoter regions increases with increasing stage of malignancy in various human cancers and often results in gene silencing (Jones, 2012). On the other hand, loss of methylation can be predictive of increased gene expression (Weigel et al, 2016). Novel technologies based on genome-wide screens for aberrant DNA methylation and epigenetic gene silencing, including the widely used Illumina 450k Beadchip arrays, have allowed identification of hundreds of genes aberrantly methylated during prostate cancer development (Borno et al, 2012; Brocks et al, 2014; Cancer Genome Atlas Research, 2015; Fraser et al, 2017; Kim et al, 2011; Kobayashi et al, 2011; Kron et al, 2009; Weischenfeldt et al, 2013). These cancer-specific epigenetic alterations have been shown to enable the development of methylation-based assays to distinguish between benign and malignant tissue and to predict the course of the disease (Haldrup et al, 2013; Jeronimo et al, 2011; Nelson et al, 2009; Yang & Park, 2012).

In recent years, machine-learning techniques became widely used in modern molecular research to build predictive models (Camacho et al, 2018). Random forest (Breiman, 2001) is an ensemble learning method based on the construction of many classification trees. Main benefits of the method are its robustness against overfitting, user-friendliness and the easy interpretation of the model (Breiman, 2001).

Our goal was to use random forest-based modelling of DNA methylation alterations to develop a classifier predicting the outcome of PCa. The tight connection of DNA methylation events with gene expression allowed us to utilize immunohistochemistry (IHC), a universally available tool in diagnostic laboratories, on tissue microarrays of thousands of clinically well annotated samples to validate ZIC2 as a prognostic protein biomarker independent of established clinico-pathological variables.

## Results

### Differential methylation analysis

To identify methylation alterations associated with PCa aggressiveness, we used a discovery cohort of 70 PCa cases (**Table 1**) with good (organ confined disease and lack of recurrence for at least 5 years) or poor prognosis (systemic presence of metastatic disease, indicated by biochemical prostate-specific antigen (PSA)-based recurrence (BCR) within 3 years and no response to local radiation therapy) for genome-wide methylation analyses using Illumina 450k arrays. The two groups showed differences in preoperative PSA levels (p=1.2*10^−6^). As a result of the selection criteria, the two sample groups are very different regarding their survival rates. In fact, patients in the poor prognosis group suffered from rapid BCR, with a median disease-free survival of only 3.8 months.

**Table 1:** Clinical characteristics of the discovery cohort.

|  | **Good prognosis<sup>a</sup>** | **Poor prognosis<sup>a</sup>** |
| --- | --- | --- |
| **n** | 35 | 35 |
| **Age** (mean ± sd) | 62.7 ± 5.6 | 65 ± 6.6 |
| **Pretreatment PSA** (ng/ml) | 6.86 ± 3.4 | 28.3 ± 22.3 |
| **Stage (path. T)** |  |  |
| pT2 | 35 | 0 |
| pT3a | 0 | 3 |
| pT3b | 0 | 31 |
| pT4 | 0 | 1 |
| **Gleason score** |  |  |
| 3+3 | 15 | 0 |
| 3+4 | 17 | 6 |
| 4+3 | 3 | 15 |
| 4+4 | 0 | 4 |
| 4+5 | 0 | 7 |
| 5+4 | 0 | 3 |

<sup>a</sup> Good prognosis defined as an organ confined disease (pT2) and lack of biochemical (PSA based) recurrence (BCR) for at least 5 years. Bad prognosis defined as systemic presence of metastatic disease, indicated by recurrence within 3 years and no response to local radiation therapy.

After adjusting for age at diagnosis and the samples’ basal, stromal and immune cell contents computed from DNA methylation data using the PEPCI package (Gerhauser et al, 2018) (**Supplementary Table 1**), we selected 402 differentially methylated CpG sites (DMS, minimum 10% absolute methylation difference, FDR adjusted p-value < 0.2) (**Figure 1A**). Of these, 302 DMS lost methylation in the bad prognosis group compared to the good prognosis group, and 100 DMS gained methylation (**Figure 2**). Hypermethylated DMS were mainly localized in CpG islands, shores and shelves, while the ones with lower methylation were mostly located in intergenic (open sea) regions (**Figure 1B**). Using chromatin state information based on ChromHMM data for normal prostate (PrEC) and prostate cancer (PC3) cell lines (Taberlay et al, 2014), we confirmed that the hypermethylated DMS were overlapping with poised promoters and repressed regions, while hypomethylated DMS showed enrichment for heterochromatic regions (**Figure 1C**). In line with these results, a GREAT-based pathway analysis showed enrichment of developmental processes among genes affected by hypermethylation (**Figure 1D**).

**Figure 1.**
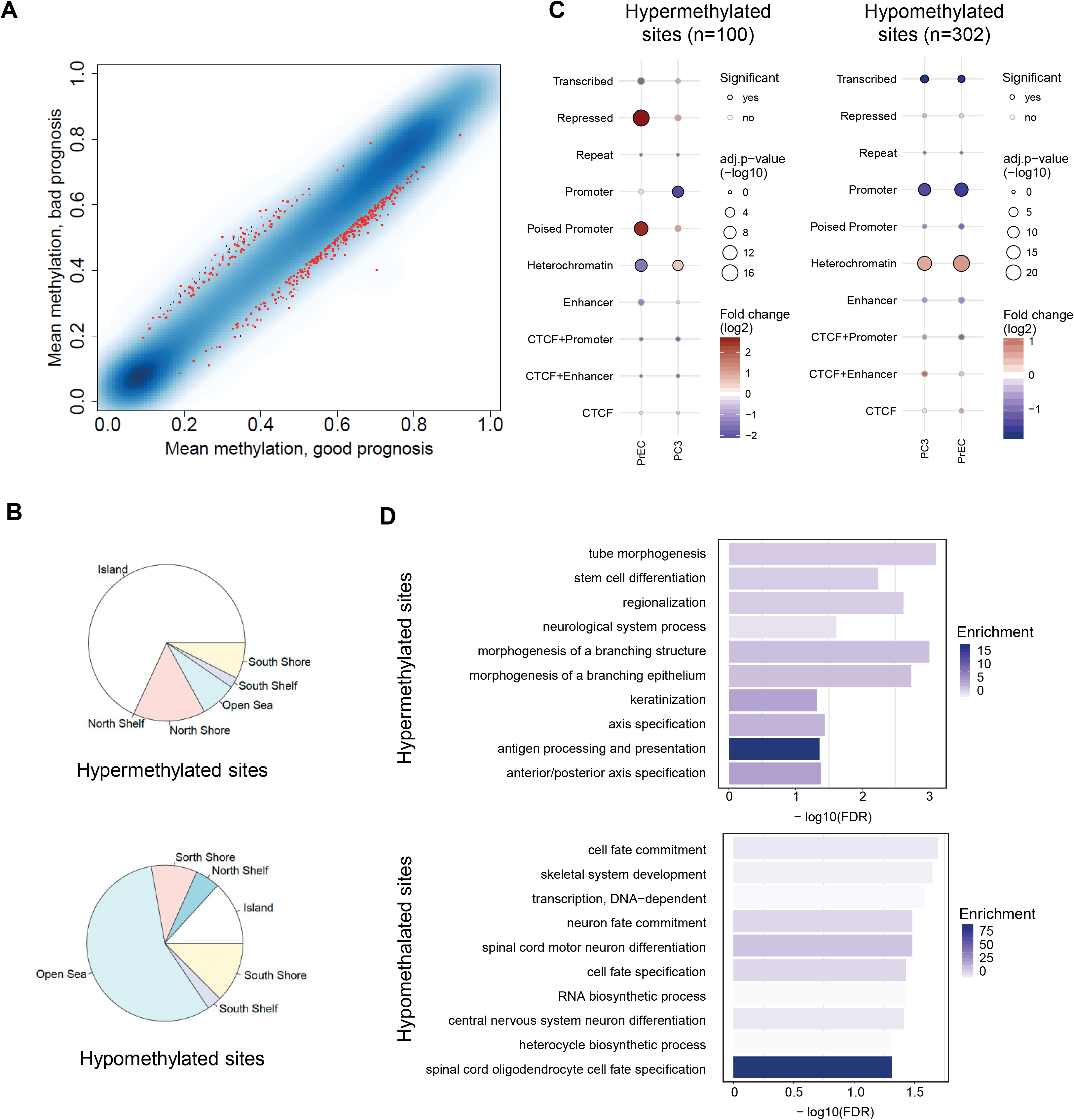
Differential methylation analysis. A. Mean methylation values of the good and the bad prognosis groups shown in a smoothed colour density representation plot. The high density parts of the plot are smoothed, only the low density parts are shown as separate points. The further away a point is from the diagonal, the more different it is in the two subgroups. Sites with FDR p value < 0.2 and absolute beta difference > 0.1 are marked in red. B. Distribution of the localization of differentially methylated CpG sites (DMS) hypermethylated (n=100, left) and hypomethylated (n=302, right) in the bad prognosis group relative to the good prognosis group, in relation to CpG islands. C. Enrichment analysis of the hypermethylated (left) and hypomethylated (right) DMS using ChromHMM data for PC3 and prostate epithelial cells (PrEC) (Taberlay et al, 2014). The size of the circles shows the significance of enrichment, while its colour represents the strength and the direction of the enrichment (red - enriched, blue - depleted). The black circle outline shows that the result is significant. D. Pathway analysis of genes associated with the hypermethylated (left) and hypomethylated (right) DMS using the GREAT tool. The shade of the blue shows the significance of the enrichment, while the bars represent the strength.

**Figure 2.**
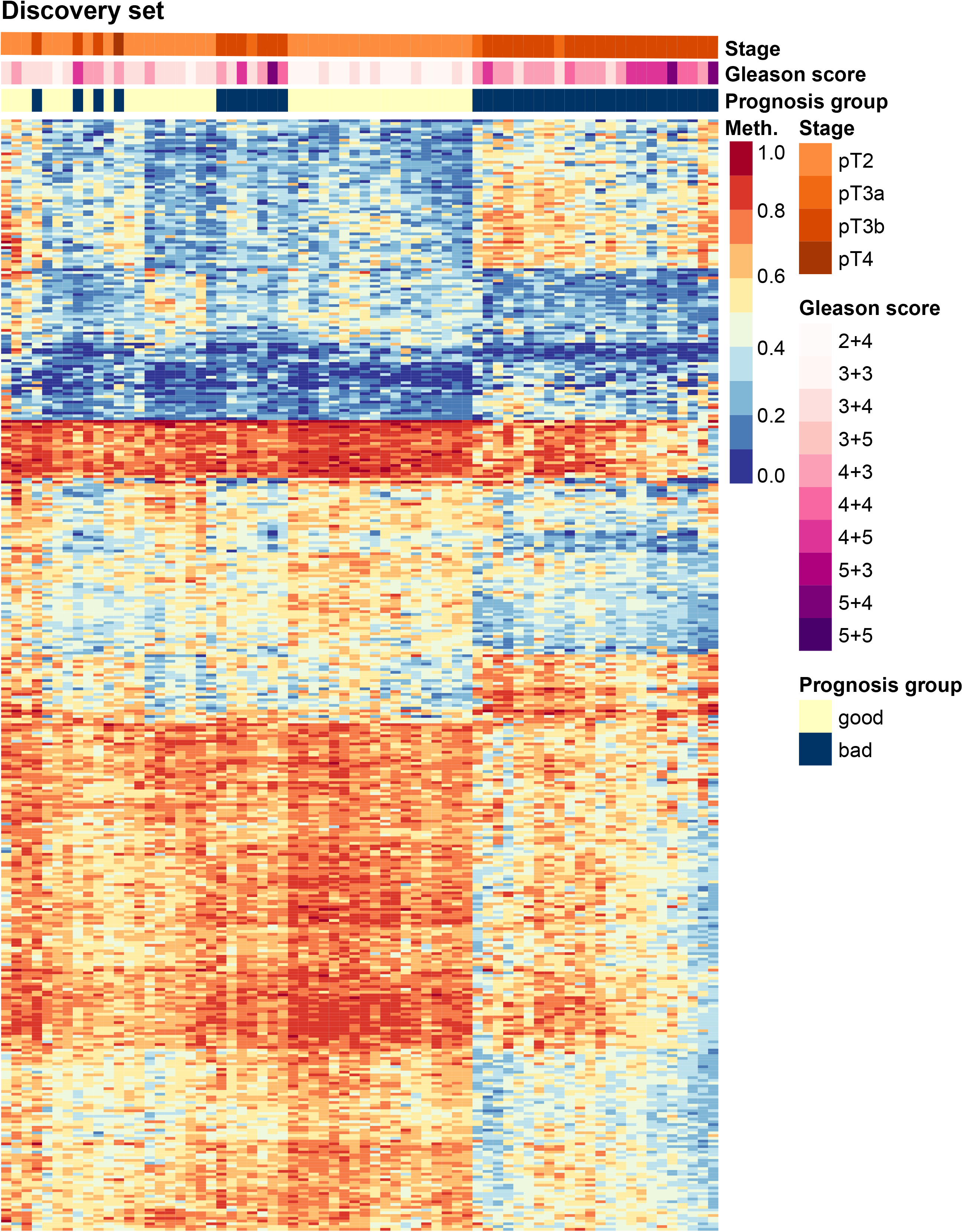
Methylation heatmap of the selected CpG sites in the discovery cohort. Each column represents a sample with either good or bad prognosis, while rows represent methylation beta values of each selected CpG sites in a range from 0 to 1.

### Random forest model

We applied random forest-based modelling to rank the selected DMS according to their discriminative power (for details see description in the Experimental section). In addition to the DMS, the Purity-Adjusted Epigenetic Prostate Cancer Index (PEPCI) of tumor aggressiveness (Gerhauser et al, 2018) was included in the model. Mean PEPCI was significantly different between the two prognosis groups of the discovery cohort (t-test p-value=0.03). Using a cut-off of 69.1 to define PEPCI-low and PEPCI-high tumors (Gerhauser et al, 2018), the aggressivity score stratified the discovery cohort according to PSA recurrence-free survival (log-rank p-value=0.045)(**Supplementary Figure 2**). For the random forest-based modelling, the discovery cohort was randomly split into a training (80% randomly selected samples) and a test set (20% randomly selected samples). The model was trained on the training set, with 10000 trees. Prediction accuracy was then measured on the test set. For variable selection, DMS were ranked based on mean decrease in accuracy of the model and mean decrease in Gini scores (Hastie et al, 2009) (complete list of CpG sites in the model, as well as importance scores in **Supplementary Table 2**). The Gini score indicates how often a random sample from the test set would be incorrectly categorized as having good or bad prognosis if the samples were randomly distributed (Hastie et al, 2009).

The random forest model showed an error of 14.81% on the training set (n=56), with better prediction for the bad prognosis subgroup. On the test set (n=14), the model showed an error rate of 18.8%, with an area under the receiver operating characteristic (ROC) curve (AUC) of 95% (**Figure 3A**). A Kaplan-Meier plot indicated excellent stratification of the subgroups of the test set predicted to have good or bad prognosis (log-rank p-value <0.0001, **Figure 3B**). We applied our model to two independent PCa cohorts for validation of the good prediction rate. We were able to validate our results using the ICGC PCa cohort of early and late onset prostate cancer (n= 222) (Gerhauser et al, 2018). The AUC for the sensitivity analysis was 77.1% and the error rate was 35.1% (**Figure 3C**). With an AUC of 99.7% and an error rate of 1.6%, the model demonstrated excellent performance when we only used a subset of the cohort based on the same selection criteria as in our discovery cohort (n=63). With the TCGA PRAD cohort (n=477, **Table 2**), we observed an AUC of 68.7% and an error rate of 43% (**Figure 3E**). With a preselected subset (n=84, **Table 2**), we achieved an AUC of 77.5% and an error rate of 28.6%. Our classifier stratified both validation cohorts according to PSA recurrence-free survival (log-rank p-value <0.0001 for both cohorts) (**Figure 3D** and **F**, and **Supplementary Figure 3**). In the ICGC dataset, our model proved to be an independent predictor of recurrence-free survival, when the Gleason score was included in the model (Cox regression p=0.011).

**Figure 3.**
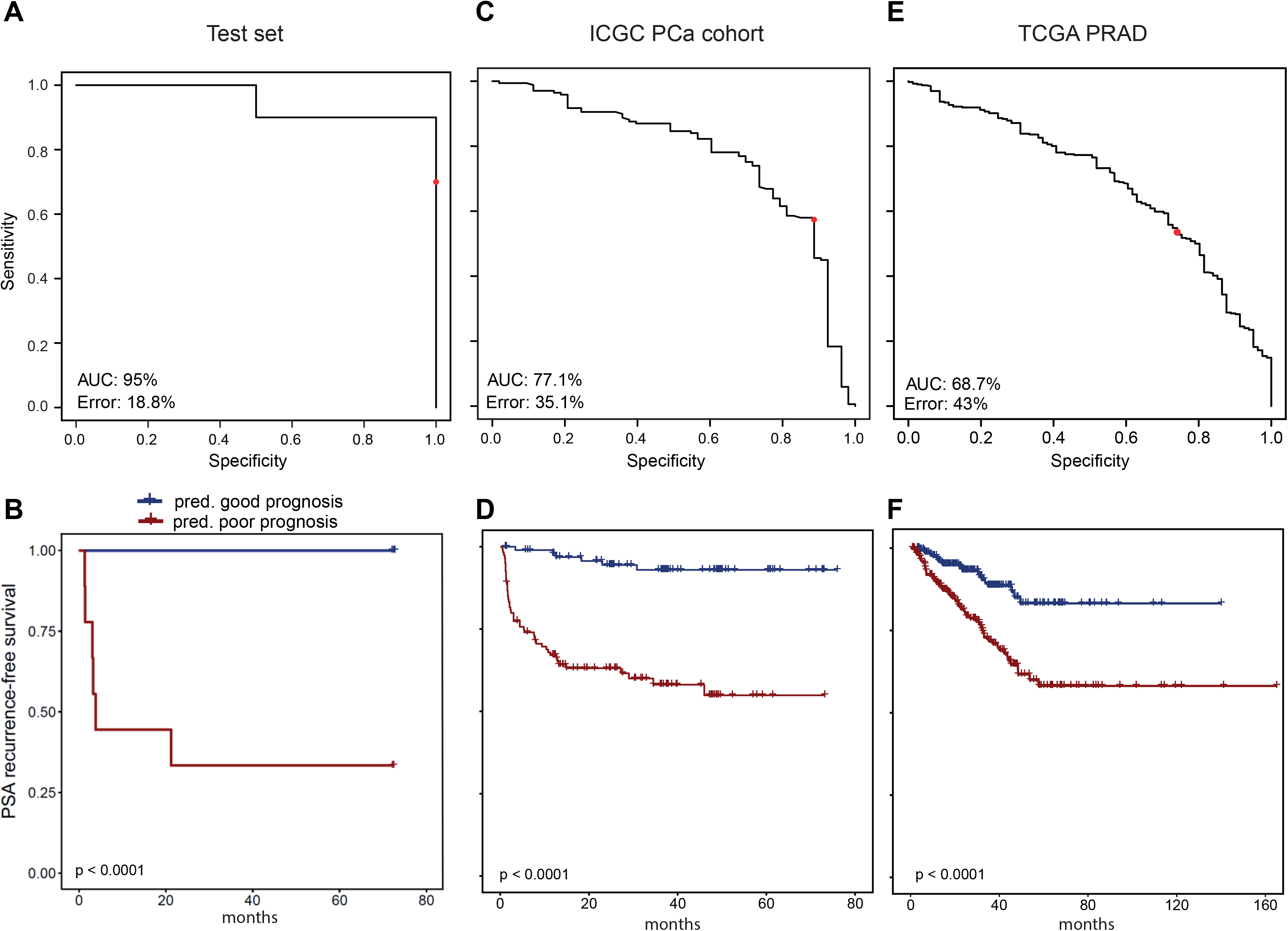
Performance analysis of the model in the test and validation datasets. A,C,E. ROC curve analysis of the model’s performance in the test set (A), ICGC PCa cohort (C) and TCGA PRAD cohort (E). The x-axis shows the model’s specificity while the y-axis shows the sensitivity. Red dot represents the performance of the model when a cut-off of 0.5 is used during classification. B,D,F. Kaplan-Meier curves using PSA recurrence-free survival as an outcome in the test set (B), in the ICGC PCa cohort (D), and in the TCGA-PRAD cohort (F), based on the predicted prognostic categories (blue: good prognosis, red: poor prognosis). p values were calculated using log-rank test.

**Table 2:**
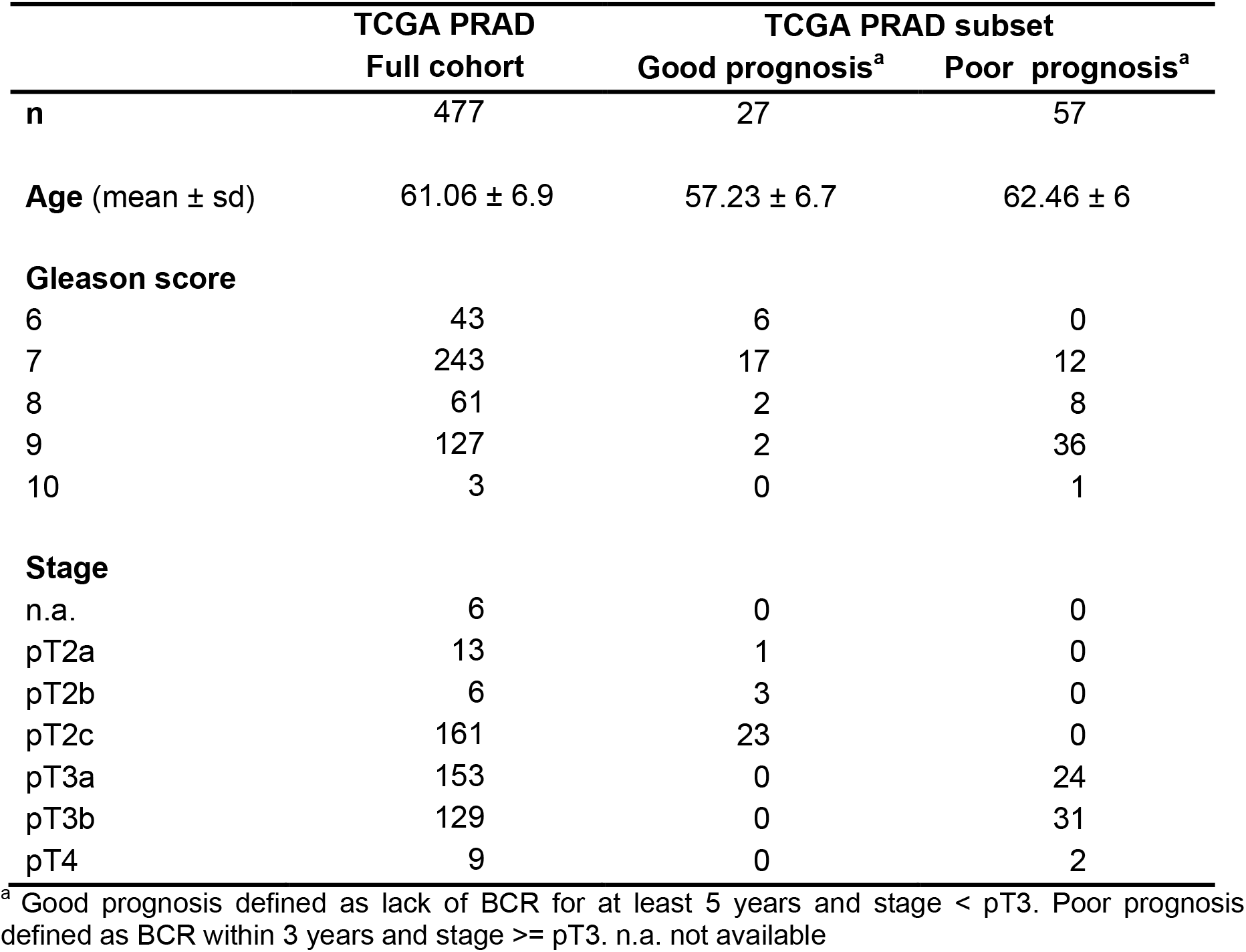
Clinical characteristics of the TCGA PRAD cohort and the preselected subset.

### Candidate selection

The relevance of the DMS included in the model is supported by previous literature. They are often located in promoter or enhancer regions of genes of importance for the aggressiveness or progression of prostate cancer. Based on their localization in regulatory regions and distance from transcription start sites (TSSs), DMS were associated to genes. The genes were ranked to select the top10 candidates for confirmatory analyses based on immunohistochemistry (IHC), as further described below (**Table 3**, with individual Kaplan-Meier curves for the candidate gene-related CpG sites and the full model in **Supplementary Figure 2**).

**Table 3.** Candidate CG probes and genes.

| CG probe names | Chr. | Location | Mean decrease in accuracy | Mean decrease in Gini | ChromHMM PC3 | ChromHMM PrEC | Genes | Distance from TSS | Mean meth. good progn. | Mean meth. poor progn. |
| --- | --- | --- | --- | --- | --- | --- | --- | --- | --- | --- |
| cg14496282 | chr7 | 544525 | 0.002256 | 0.223239 | Enhancer | Enhancer | PDGFA | 14505 | 0.509 | 0.631 |
| cg07043604 | chr8 | 48739256 | 0.001495 | 0.159048 | Enhancer | Enhancer | PRKDC | 133486 | 0.662 | 0.554 |
| cg16876647 | chr2 | 43451842 | 0.002281 | 0.15899 | Promoter | Promoter | ZFP36L2 | 1905 | 0.703 | 0.401 |
| cg12092201 | chr2 | 43451775 | 0.001586 | 0.122542 | Promoter | Promoter | ZFP36L2 | 1972 | 0.473 | 0.285 |
| cg04211581 | chr6 | 152011656 | 0.000859 | 0.117083 | CTCF | Heterochromatin | ESR1 | 26 | 0.458 | 0.317 |
| cg24690071 | chr13 | 100635352 | 0.001171 | 0.116259 | Poised_Promoter | Repressed | ZIC2 | 1327 | 0.483 | 0.322 |
| cg00874055 | chr1 | 236306673 | 0.001063 | 0.115871 | Promoter | Promoter | GPR137B | 842 | 0.487 | 0.342 |
| cg05131488 | chr7 | 14031822 | 0.000886 | 0.104012 | Enhancer | Heterochromatin | ETV1 | -2252 | 0.423 | 0.551 |
| cg12658720 | chr17 | 59488443 | 0.00112 | 0.096522 | CTCF+Enhancer | Promoter | TBX2 | 11187 | 0.558 | 0.446 |
| cg12658720 | chr17 | 59488443 | 0.00112 | 0.096522 | CTCF+Enhancer | Promoter | TBX4 | -45363 | 0.558 | 0.446 |
| cg11462315 | chr2 | 43455048 | 0.000689 | 0.088083 | CTCF+Promoter | Promoter | ZFP36L2 | -1301 | 0.271 | 0.162 |
| cg21940313 | chr17 | 41620911 | 0.000665 | 0.079846 | Promoter | Enhancer | ETV4 | 2888 | 0.502 | 0.401 |
| cg10981580 | chr1 | 236306767 | 0.000794 | 0.07559 | Promoter | Promoter | GPR137B | 936 | 0.418 | 0.282 |
| cg08738430 | chr2 | 43452561 | 0.000608 | 0.074411 | Promoter | Promoter | ZFP36L2 | 1186 | 0.264 | 0.110 |

Among the top candidate DMS, five sites affected the promoter region of zinc-finger proteins. Four sites were located in the promoter of *ZFP36L2* (cg16876647, cg12092201, cg11462315, cg08738430), while a poised promoter of *ZIC2* was affected by one CpG site (cg24690071). *ZFP36L2* encodes a CCH-type zinc finger protein, which is regulated by the cell-cycle, might play a role in DNA damage response (Noguchi et al, 2018), and inhibits cell proliferation (Liu et al, 2018; Suk et al, 2018). In PCa, ZFP36L2 upregulation was associated with the transcription factor Runx2 and poor prognosis (Baniwal et al, 2010). ZIC2 belongs to a family of transcription factors involved in neuro-ectodermal development. Elevated ZIC2 mRNA expression was described in high Gleason prostate cancer (Hoogland et al, 2016).

Cg05131488 and cg21940313 are located in the enhancer and promoter regions of *ETV1* and *ETV4*. These are both members of the erythroblast transformation-specific (ETS) transcription factor family along with *ERG* and commonly affected by gene fusions in PCa (Tomlins et al, 2006; Tomlins et al, 2005). Their overexpression showed oncogenic effects with slightly overlapping functions (Mesquita et al, 2015).

Cg04211581 is located only 26 bps from the TSS of *ESR1*. ESR1 encodes estrogen receptor alpha (ERα), the role of which was proposed in PCA, however, is still controversial (Yeh et al, 2014).

*TBX2* and *TBX4* came up as candidate genes, since cg12658720 lies in an enhancer region between them. As members of the T-box (TBX) protein family, TBX proteins, mainly *TBX2* and *TBX3* play important oncogenic role in several types of cancer (Chang et al, 2016).

Platelet-derived growth factor (PDGF) acts as an oncogene promoting tumor growth and survival in prostate cancer, as well as in other cancer types. It is commonly overexpressed in prostate cancer, therefore represents a potential therapeutic target (Heldin, 2013).

A highly predictive CpG site, cg07043604, is located in an enhancer region around 130kb away from *PRKDC*, which encodes the catalytic subunit of DNA-dependent protein kinase (DNA-PKcs). Previous research identified DNA-PKcs as a master regulator of pro-metastatic signaling and showed that its expression is strongly correlated with survival (Goodwin et al, 2015).

The influence of the observed methylation changes of these candidates DMS on gene or protein expression and the impact on prostate carcinogenesis needs to be experimentally confirmed in mechanistic chromatin conformation and gain- and loss-of-function studies.

### Candidate validation

ZIC2 was one of the candidate genes for which a suitable antibody for IHC was available. ZIC2 expression was analysed by immunohistochemistry on a tissue microarray (TMA) containing more than 12000 prostate cancer specimens (**Table 4**). Results were compared with tumour phenotype, BCR, ETS-related gene (ERG) status and other recurrent genomic alterations. ZIC2 expression was detectable and considered to be strong in 23.3% of cases and was absent in majority of the tumours (76.7%) (**Figure 4A**, **Table 4**). Loss of ZIC2 protein expression was associated with ERG-fusion positivity (p< 0.0001) (**Figure 4B**). Loss of ZIC2 expression was also linked to Gleason grade, advanced pathological tumour (pT) stage, lymph node metastasis and higher preoperative PSA levels in all cancers (p<0.0001, each) and in the subset of ERG-fusion negative tumours (**Table 4**, data not shown). These associations were either weaker or absent in ERG-fusion positive cancers (data not shown). Within ERG-fusion negative cancers, ZIC2 expression was also strongly associated with 6q15 and 5q21 deletions (p<0.001) (**Figure 4C**). Loss of ZIC2 expression was associated with adverse outcome and correlated with significantly shorter time to biochemical recurrence in all cancers, independent of ERG and PTEN (**Figure 4D**). The prognostic value of ZIC2 proved to be independent from established clinico-pathological variables including Gleason, stage, nodal stage and PSA. Overall, ZIC2 was identified as an excellent marker and might provide clinically useful predictive information by identification of aggressive prostate cancer subsets.

**Table 4:**
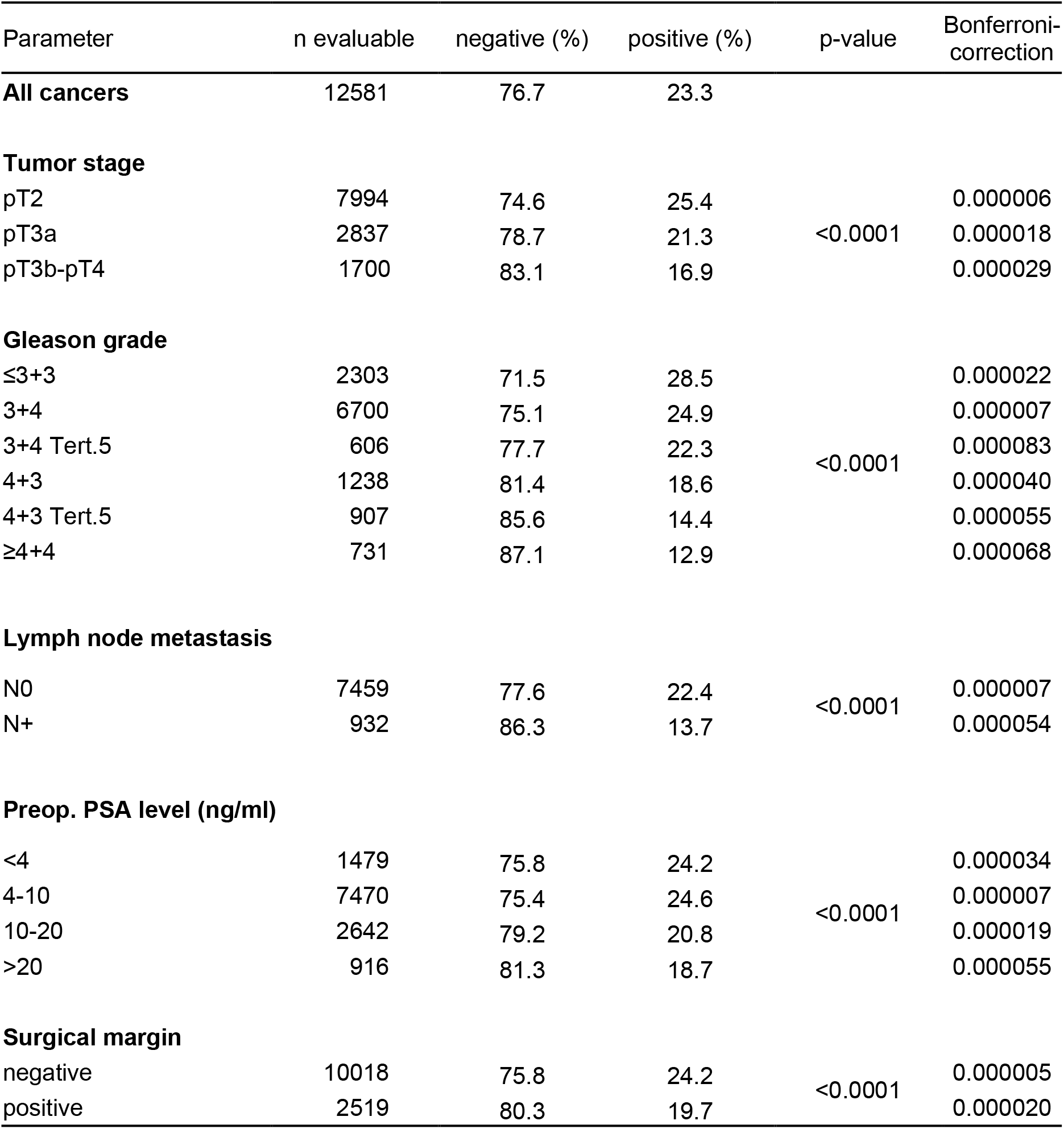
Association between ZIC2 immunostaining results and prostate cancer phenotype in all cancers.

| Parameter | n evaluable | negative (%) | positive (%) | p-value | Bonferroni-correction |
| --- | --- | --- | --- | --- | --- |
| **All cancers** | 12581 | 76.7 | 23.3 |  |  |
| **Tumor stage** |  |  |  |  |  |
| pT2 | 7994 | 74.6 | 25.4 | <0.0001 | 0.000006 |
| pT3a | 2837 | 78.7 | 21.3 |  | 0.000018 |
| pT3b-pT4 | 1700 | 83.1 | 16.9 |  | 0.000029 |
| **Gleason grade** |  |  |  |  |  |
| ≤3+3 | 2303 | 71.5 | 28.5 | <0.0001 | 0.000022 |
| 3+4 | 6700 | 75.1 | 24.9 |  | 0.000007 |
| 3+4 Tert.5 | 606 | 77.7 | 22.3 |  | 0.000083 |
| 4+3 | 1238 | 81.4 | 18.6 |  | 0.000040 |
| 4+3 Tert.5 | 907 | 85.6 | 14.4 |  | 0.000055 |
| ≥4+4 | 731 | 87.1 | 12.9 |  | 0.000068 |
| **Lymph node metastasis** |  |  |  |  |  |
| N0 | 7459 | 77.6 | 22.4 | <0.0001 | 0.000007 |
| N+ | 932 | 86.3 | 13.7 |  | 0.000054 |
| **Preop. PSA level (ng/ml)** |  |  |  |  |  |
| <4 | 1479 | 75.8 | 24.2 | <0.0001 | 0.000034 |
| 4-10 | 7470 | 75.4 | 24.6 |  | 0.000007 |
| 10-20 | 2642 | 79.2 | 20.8 |  | 0.000019 |
| >20 | 916 | 81.3 | 18.7 |  | 0.000055 |
| **Surgical margin** |  |  |  |  |  |
| negative | 10018 | 75.8 | 24.2 | <0.0001 | 0.000005 |
| positive | 2519 | 80.3 | 19.7 |  | 0.000020 |

**Figure 4.**
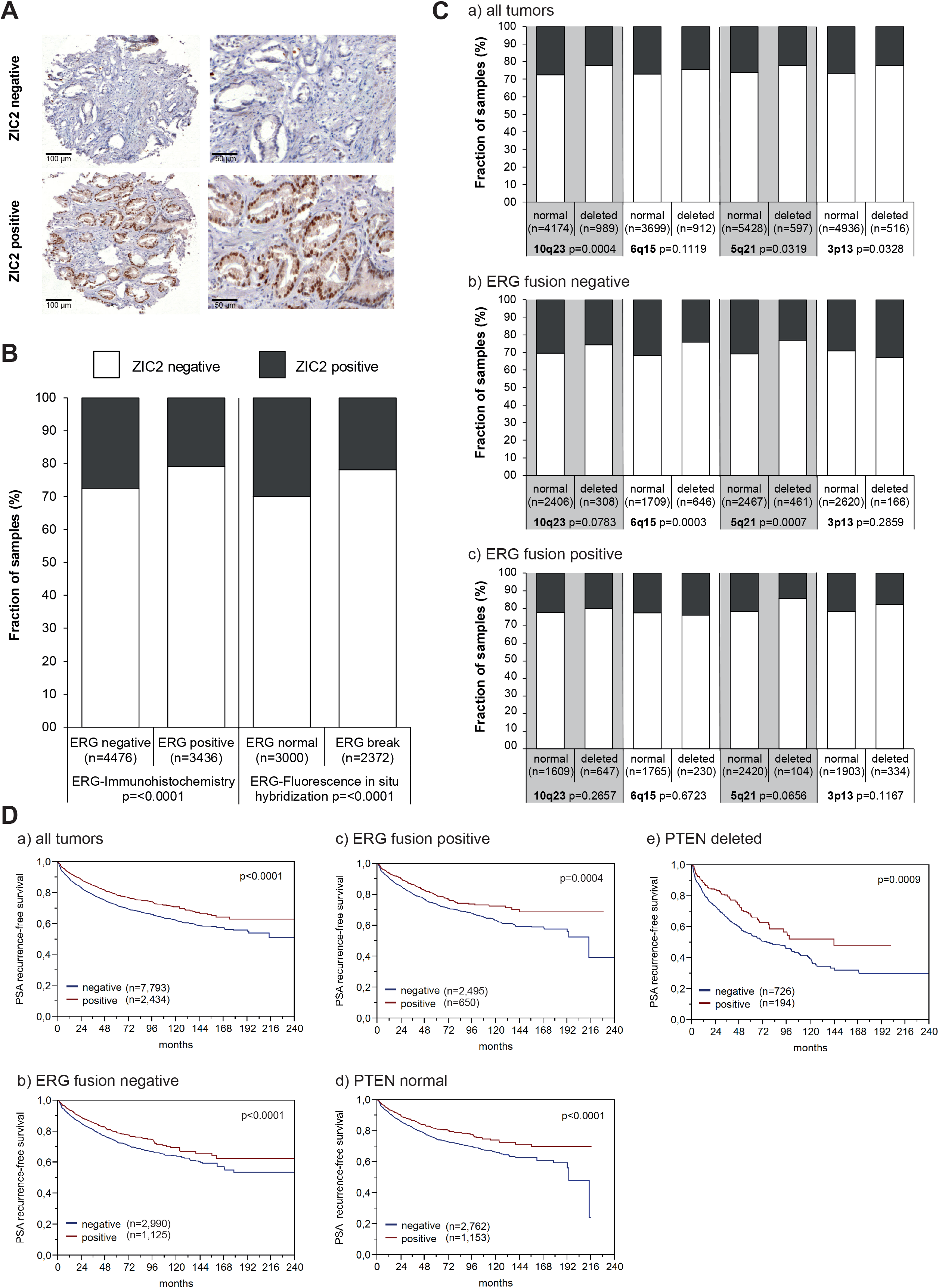
ZIC2 immunostaining in >12000 micro-arrayed PCa cases. A. Examples of negative (no nuclear staining, upper panels) and strong staining (lower panels). B. Association between ZIC2 immunostaining results and the ERG-status determined by IHC and FISH analysis. C. Association between ZIC2 immunostaining and deletions of 10q23 (PTEN), 6q15 (MAP3K7), 5q21 (CHD1), and 3p13 (FOXP1) for all cancers (a), ERG fusion negative (b) and ERG fusion positive subset (c) according to ERG-IHC analysis. D. Kaplan-Meier curves for the relationship of ZIC2 immunostaining with PSA recurrence-free survival in all cancers (a), in ERG fusion negative cancers (b), in ERG fusion positive cancers (c), in PTEN normal cancers (d), and in PTEN deleted cancers (e). Log-rank p-values.

## Discussion

In the present study we have identified methylation differences related to PCa prognosis, and subsequently showed that methylation-based prediction of PCa prognosis using random forest-based modelling is feasible with high accuracy.

PCa is the most prevalent cancer among men in Germany. The 5-year survival rate is 91% (<u>German Cancer Registry</u>), since in many cases the cancer remains indolent, with no need of aggressive treatment. However, some patients develop an aggressive from of the disease. Therefore, biomarkers predicting the prognosis of PCa are needed for an efficient clinical management of this disease (Prensner et al, 2012).

DNA methylation is an excellent source for biomarker development, since it is a stable modification and can be quantitatively determined in clinical samples with high throughput and precision and relatively low cost (Claus et al, 2012). Previous studies trying to establish a methylation-based classifier for prostate cancer mostly used a preselected set of genes (Ahmad et al, 2016; Litovkin et al, 2015), or used high Gleason score as an outcome (Bhasin et al, 2015; Geybels et al, 2016). Here we are presenting a genome-wide approach, with PSA recurrence-free survival as an endpoint, to achieve the strongest relevance for clinical decisions. One limitation of our study is the use of the Illumina 450k platform for biomarker selection, which limits methylation analyses to preselected CpG sites on the 450k array (enriched for CpG islands and flanking regions, bioinformatically predicted enhancers, DNase I hypersensitive sites, and validated differentially methylated regions (Stirzaker et al, 2014)). Future studies using whole genome bisulfite sequencing (WGBS) of all >29 Mio CpG sites in the human genome will allow identification of additional biomarkers.

Our discovery cohort consisted of 70 patients, 35 with good and 35 with bad prognosis. After cell type adjustments, our cut-off criteria for selection of differentially methylated CpG sites were mean beta difference > 0.1 and an FDR-adjusted p-value < 0.2. Altogether, 402 DMS and the PEPCI score for tumour aggressiveness (Gerhauser et al, 2018) were included in the prediction model. Our random forest-based model showed good performance with the discovery cohort. We were able to validate our results using the ICGC PCa cohort of early and late prostate cancer (AUC 77.1%), with slightly worse performance using the TCGA PRAD dataset (AUC 68.7%). Different reasons might contribute to the lower performance with the TCGA-PRAD cohort, such as possibly different definitions of PSA recurrence-free survival, and the generally high Gleason score and high tumor stage of the TCGA patients. Other genome-wide studies have faced similar problems using TCGA as a validation set (Bhasin et al, 2015; Mundbjerg et al, 2017). Nevertheless, for both ICGC and TCGA validation cohorts, the resulting prognostic subgroups had highly significantly different survival rates.

A recent proteomics-based biomarker study of curable prostate cancer reported a stronger link of DNA methylation status to protein than mRNA abundance (Sinha et al, 2019). In line with these findings, we performed a validation of the clinical impact of ZIC2 as one of the candidate genes on more than 12000 micro-arrayed PCa cases. The *zinc finger of the cerebellum* (ZIC) family of genes consists of five human homologues ZIC1–5 (Ali et al, 2012). ZIC family members inhibit TCF4/β-Catenin and interact with GLI signalling (Ishiguro et al, 2018). In addition, an oncogenic role of ZIC family members has been reported. ZIC1, −2 and −5 are over-expressed in meningioma, while ZIC4 is over-expressed in medulloblastomas (Aruga et al, 2010). Methylation of ZIC4 in bladder cancer is predictive for progression (Beukers et al, 2015). Promoter methylation of ZIC1 is associated with gastric cancer (Schneider et al, 2013; Wang et al, 2009). ZIC2 is related to the sonic hedgehog pathway. Its oncogenic role was described in epithelial ovarian cancer (Marchini et al, 2012), hepatocellular carcinoma (Lu et al, 2017) and pancreatic cancer (Inaguma et al, 2015). High expression of ZIC2 was associated with survival in oral squamous carcinoma (Sakuma et al, 2010). Our IHC validation indicated a particularly strong adverse prognostic value of ZIC2 expression loss. ZIC2 analysis appears to be of high value for distinguishing between patients with more or less aggressive forms of the disease and may be useful to select patients for active surveillance. Our results may thus help to reduce the number of unnecessary prostatectomies.

In summary, we present a candidate selection of cancer progression-related CpG sites, as well as a classification model to predict aggressive behaviour of PCa. This model, with further tuning, might help in decision making related to the treatment of prostate cancer patients. The effect of candidate CpG site methylation on gene expression helps to pinpoint further genes, which play an important role in prostate cancer. Ranking of the selected CpG sites and associated genes allowed selection of candidates for validation by IHC. We identified loss of ZIC2 expression as a promising prognostic biomarker for PCa.

## Material and Methods

### Study population

In order to build a classifier that predicts the patients’ outcome the best, a highly selected group of patients was included in the study. Sample selection was based on the following criteria: good prognosis indicated by presence of organ confined disease (pT2) and lack of biochemical PSA-based recurrence for at least 5 years. In contrast, bad prognosis is defined as systemic presence of metastatic disease, indicated by BCR within 3 years and no response to local radiation therapy. Initially, 84 patients were selected. All PCa samples had been collected between 1992 and 2012 from routine diagnosis leftover tissues, the usage of which for research purposes is legally covered by §12 of the Hamburgisches Krankenhausgesetz. The local ethics committee approved usage of these tissues for TMA manufacturing (WF-049/09, Ethik-Kommision der Ärztekammer Hamburg).

A pathologist selected FFPE tissue blocks containing tumour-rich areas (≥70% tumour cells) for analysis. Three tissue punches (0.6 mm × 3 mm) were taken of each tissue block and genomic DNA was isolated using the AllPrep® DNA/RNA FFPE kit (Qiagen). DNA was submitted to the DKFZ Genome and Proteome core facility for Illumina 450k Methylation analyses. After removing samples and DNA methylation profiles with low quality, the study included 35 patients with good and 35 patients with bad prognosis (**Supplementary Table 1**).

### Validation datasets

The ICGC PCa cohort has been described earlier (Gerhauser et al, 2018). Clinical information for the TCGA-PRAD cohort was downloaded from cBioPortal in June 2018 (**Table 2**). For the subcohorts, patients were selected as good prognosis patients by lack of BCR within 5 years and a disease stage pT2, and as poor prognosis patients when suffering from BCR within 3 years and having a stage pT3 or pT4.

### DNA methylation processing

DNA methylation was assessed using Illumina HumanMethylation450 Array. The methylation data was processed using the RnBeads R package (Assenov et al, 2014). Probes with SNPs (dbSNP 144) overlapping with the C nucleotide of the CG site and having MAF>0.01 (28722 probes) were excluded. Probes with high likelihood of false hybridization (28736 probes, as defined in RnBeads) were also removed. Quality filtering was performed using the Greedycut algorithm, which removed 21040 probes and 11 samples. Additional 969 non-CpG probes and 9229 probes located on the sex chromosomes were removed. No normalization or background correction was used. Methylation data for the discovery cohort has been uploaded to GEO under accession No. GSE127985.

During the analysis, a batch effect was observed between data from fresh frozen (TCGA PRAD and ICGC PCa cohort) and formalin-fixed tissue (discovery cohort). In order to have a generalizable model, we avoided shifting the beta values as would happen with batch correction methods. Instead, we used principal component analysis (PCA) on the top 10000 most variable CpG sites to identify the probes affected by this effect. This was done using two independent datasets containing formalin-fixed (Brocks et al, 2014) or fresh frozen tissue (Fraser et al, 2017), and removed the top 5000 sites captured by PC2, the main principal component affected by the sample type. The PEPCI score and the basal, stromal, luminal, T-luminal and immune cell composition were estimated using the PEPCI R package (Gerhauser et al, 2018). Linear models of the limma package (Ritchie et al, 2015) were applied to identify differentially methylated probes after adjustments for age, basal, stromal and immune cell content. CpG sites with FDR-adjusted p-values < 0.2 and mean methylation difference > 0.1 (10%) were used to build the model. Enrichment analysis of the significantly methylated sites, promoters and genes were performed with EpiAnnotator (Pageaud et al, 2018). Annotation of the most important CpG sites of the random forest model was done using the GREAT tool (McLean et al, 2010).

### Random forest classifier

A Random forest-based classifier was built using the randomForest R package, which is based on the algorithm of Breiman and Cutler (Breiman, 2001). Random forest is a learning method that constructs numerous decision trees and outputs the classes (in case of classification) of the individual trees. The predicted class of the input instance will be decided upon majority vote. The schematic principle of the random forest can be seen in **Supplementary Figure 4**.

The random forest uses out-of-bag (OOB) error to measure the performance of the model on the training set. In a nutshell, during the algorithm, each tree is built on a bootstrap training set, which is selected from the discovery cohort with replacement. The bootstrap training set is about two-thirds of the discovery cohort. Classification of the instances left out (OOB samples) is used to estimate a generalization error (called OOB error). The OOB error will give an unbiased estimate of the current classification error, while the bagging method will decrease the chance of overfitting.

Two variable importance scores are used in random forest. The mean decrease in accuracy reflects a variable importance measure to assess the prediction strength of each predictor variable. When a tree is grown, the OOB samples are used to calculate the error rate. Then, the values of a given predictor variable are randomly permuted and the error rate is calculated again. The decrease in accuracy caused by the permutation is averaged over all trees. The mean decrease in Gini score gives the improvement in the split-criterion at each split in each tree (Hastie et al, 2009).

Twenty different models were trained as follows: data was randomly split into training (80%) and test (20%) set. The model was trained on the training set, with 10000 trees and 19 variables to select randomly for each tree. Prediction accuracy was measured on the test set. The results were collected and the best performing model was selected. This model was then optimized for the number of variables selected for each tree. For variable selection, CpG sites were ranked based on mean decrease in accuracy and mean decrease in Gini scores (Hastie et al, 2009).

Validation of the classifier was performed using the TCGA-PRAD and the ICGC PCa cohort of early and late prostate cancer and evaluated with ROC curve analysis, using the ROCR R package (Sing et al, 2005). Performance of the prediction was measured with ROC curve analysis and Kaplan-Meier curves for the validation datasets and individual candidate CpG sites.

### Candidate selection

Based on our model, the top-rated candidates underwent a further selection to identify the ones with the highest possibility to perform well as a protein expression-based biomarker. First, we used the GREAT tool and the gene annotation of the Illumina 450k methylation array to identify the possible gene-CpG site associations, by selecting the genes closest to the sites. The CpG sites in close vicinity to genes (± 2 kb) were selected, while among the sites further away from genes, sites were included if they were located in enhancer regions based on ChromHMM data of PrEC prostate epithelial cells and the PC3 prostate cancer cell line (Taberlay et al, 2014). Finally, a literature search was conducted to identify the most relevant genes.

### Validation of candidate genes by immunohistochemistry (IHC)

#### Patients

Radical prostatectomy specimens were available from 17,747 patients undergoing surgery between 1992 and 2017 at the Department of Urology and the Martini Clinics at the University Medical Centre Hamburg-Eppendorf (**Supplementary Table 3**). All prostate specimens were analysed according to a standard procedure, including complete embedding of the entire prostate for histological analysis (Schlomm et al, 2008). Histo-pathological data was retrieved from the patient files, including tumour stage, Gleason grade, nodal stage and resection margin status. Follow-up data were available for a total of 14464 patients with a median follow-up of 48 months (range: 1 to 241 months; **Supplementary Table 3**). PSA values were measured in regular intervals following surgery and PSA recurrence was defined as the measurement of a postoperative PSA of ≥0.2 ng/ml and increasing. The TMA manufacturing process was described earlier in detail (Kononen et al, 1998). In short, one 0.6 mm core was taken from a tumour-containing tissue block from each patient. The molecular database attached to this TMA contained results on ERG expression in 10711 (Weischenfeldt et al, 2013), *ERG* break apart FISH analysis in 7122 (expanded from (Minner et al, 2011)), deletion status of 5q21 (*CHD1*) in 7932 (expanded from (Burkhardt et al, 2013)), 6q15 (*MAP3K7*) in 6069 (expanded from (Kluth et al, 2013)), 10q23 (*PTEN*) in 6704 (expanded from (Krohn et al, 2012)) and 3p13 (*FOXP1*) in 7081 (expanded from (Krohn et al, 2013)) cancers.

#### Immunohistochemistry

Freshly cut TMA sections were immunostained on one day and in one experiment. Slides were deparaffinised and exposed to heat-induced antigen retrieval for 5 minutes in an autoclave at 121 °C in pH 7.8 Tris-EDTA-Citrate buffer. Primary antibody specific for ZIC2 (antibodies online, ABIN2776475) was applied at 37 °C for 60 minutes. Bound antibody was then visualized using the EnVision Kit (Dako, Glostrup, Denmark) according to the manufacturer’s directions. ZIC2 staining intensity was assessed as negative or positive.

#### Statistics

Statistical calculations were performed with JPM 12 software (SAS Institute Inc., NC, USA). Contingency tables and the chi²-test were performed to search for associations between molecular parameters and tumour phenotype. Survival curves were calculated according to Kaplan-Meier. The log-rank test was applied to detect significant survival differences between groups. Cox proportional hazards regression analysis was performed to test the statistical independence and significance between pathological, molecular and clinical variables. Separate multivariate analyses were performed using different sets of parameters available either before or after prostatectomy.

## Acknowledgements

We thank the patients and families who contributed to this study. The study was funded by the Sander Stiftung (no. 2015.010.1). The authors acknowledge the DKFZ Genomics and Proteomics Core Facilities for excellent service and Karin Klimo and Marion Bähr (DKFZ, Division Cancer Epigenomics) for excellent technical assistance.

## Supplementary Figures

**Supplementary Figure 1:**
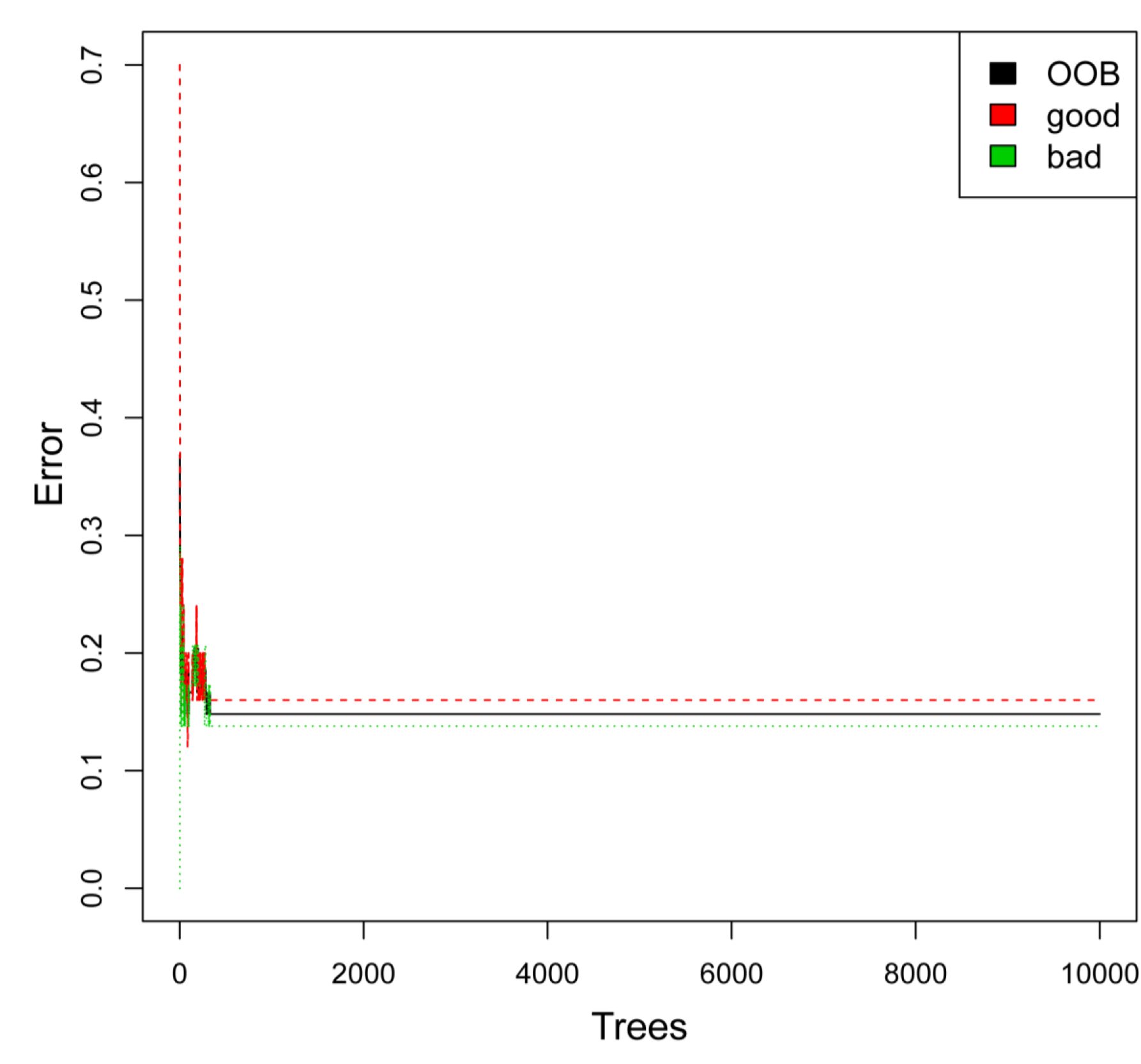
Performance of the random forest model. The plot shows the performance of the random forest model as a function of the trees built in the model, using the generalized OOB (black) and classification error for the good (red) and bad (green) prognosis groups.

**Supplementary Figure 2:**
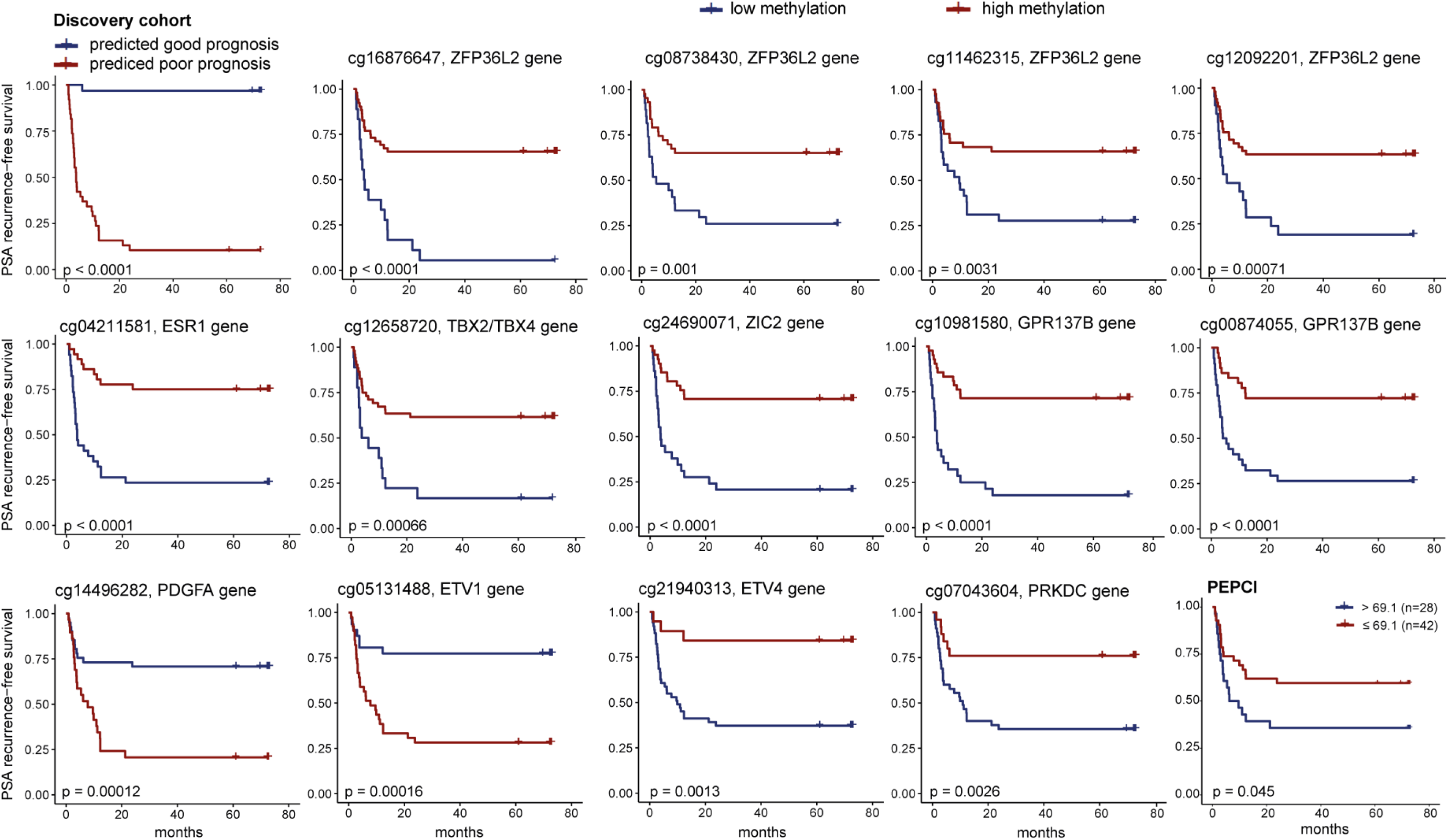
Individual Kaplan-Meier curves. Predictive power for PSA recurrence–free survival of the full classifier, PEPCI (cut-off as described previously (Gerhauser et al, 2018)), and individual candidate CpG sites associated with the top10 selected genes with the discovery cohort (n=70). p values from log-rank test. Red: high methylation (above median), blue: low methylation (below median).

**Supplementary Figure 3:**
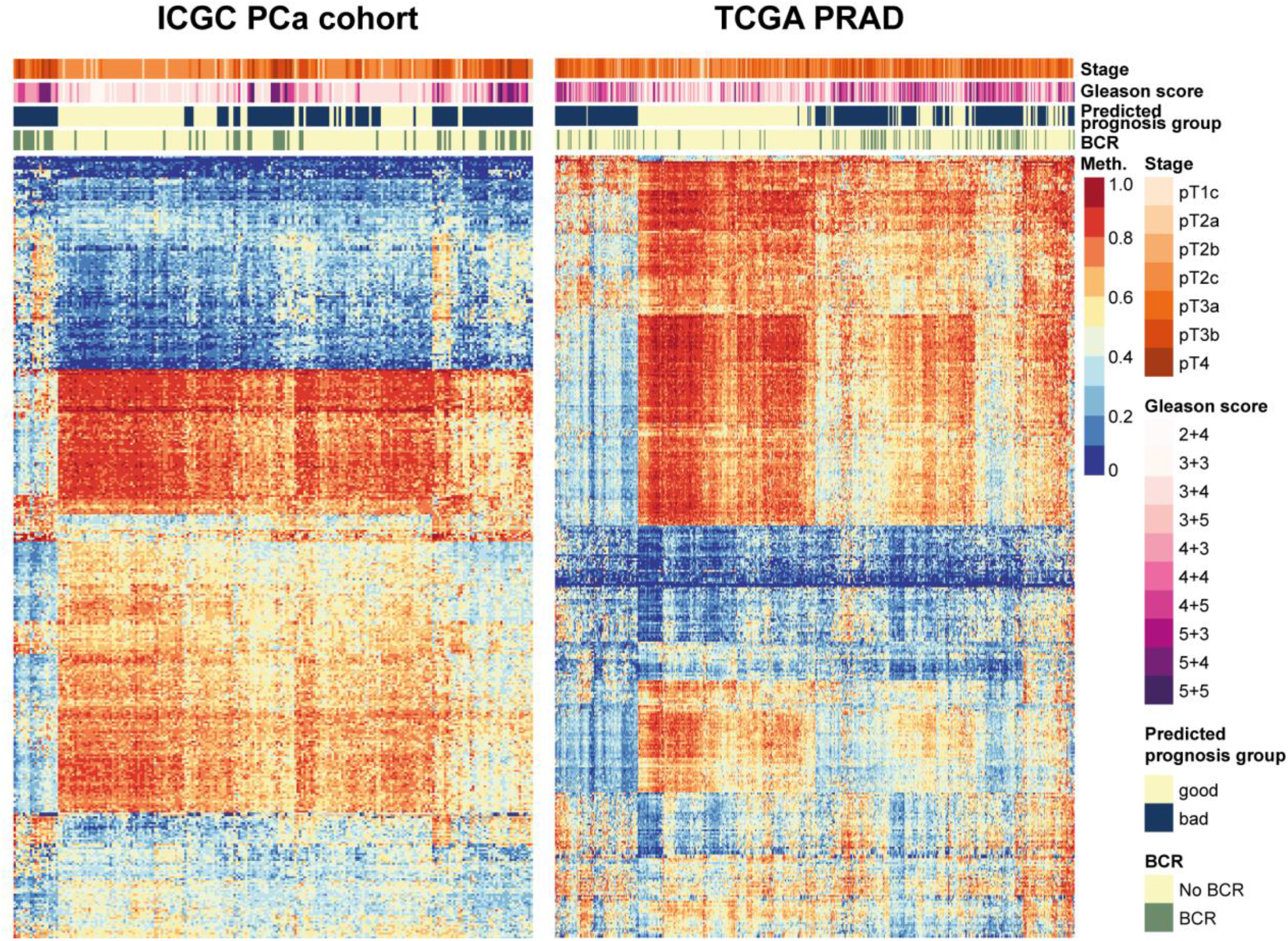
Heatmap of the selected CpG sites in the ICGC PCa (left) and TCGA PRAD (right) validation datasets. Each column represents a sample with predicted good or bad prognosis, while rows reflect on each selected CpG sites. Low and high methylation beta levels in a range from 0 to 1 are shown on a blue and red colorscale. BCR: PSA-based biochemical recurrence.

**Supplementary Figure 4.**
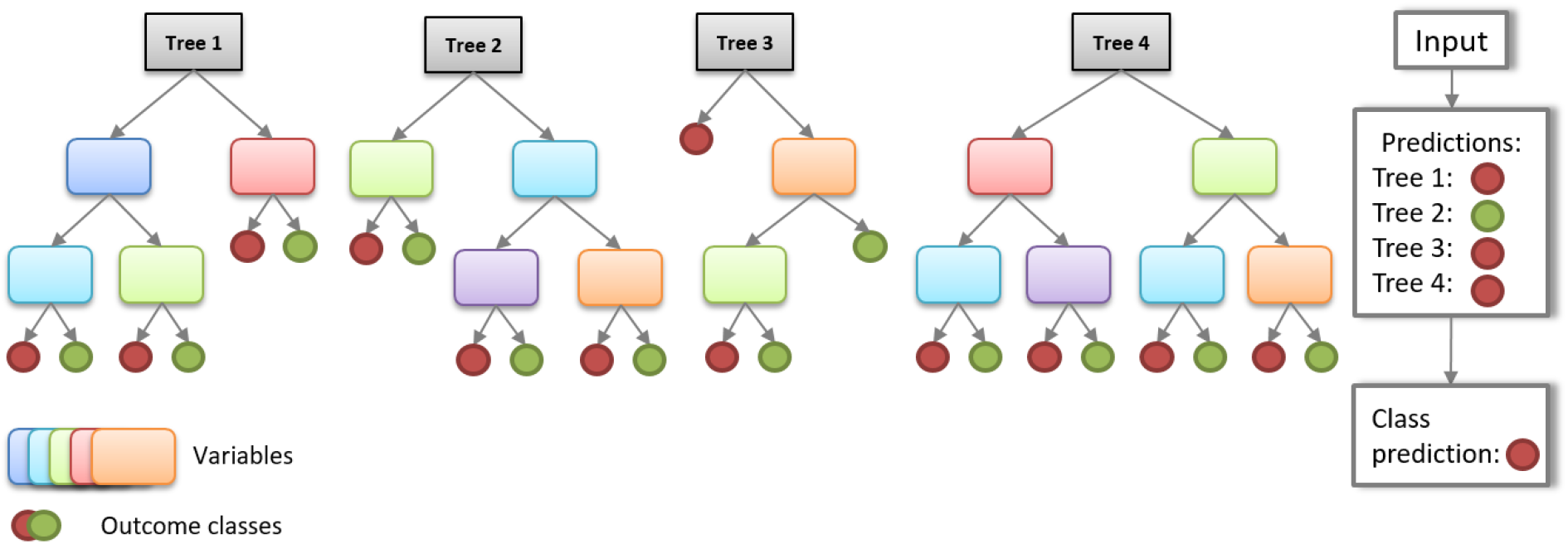
Schematic representation of the random forest model.

